# *Distal-less* and *spalt* are distal organisers of pierid wing patterns

**DOI:** 10.1101/2021.09.02.458688

**Authors:** Jocelyn Liang Qi Wee, Tirtha Das Banerjee, Anupama Prakash, Kwi Shan Seah, Antónia Monteiro

**Author notes:** Corresponding authors: Jocelyn Liang Qi Wee, Antónia Monteiro.

## Abstract

Two genes, *Distal-less (Dll)* and *spalt (sal)*, are known to be involved in establishing nymphalid butterfly wing patterns. They function in several ways: in the differentiation of the eyespot’s central signaling cells, or foci; in the differentiation of the surrounding black disc; in overall scale melanisation (*Dll*); and in elaborating marginal patterns, such as parafocal elements. However, little is known about the functions of these genes in the development of wing patterns in other butterfly families. Here, we study the expression and function of *Dll* and *sal* in the development of spots and other melanic wing patterns of the Indian cabbage white, *Pieris canidia*, a pierid butterfly. In *P. canidia*, both Dll and Sal proteins are expressed in the scale-building cells at the wing tips, in chevron patterns along the pupal wing margins, and in areas of future scale melanisation. Additionally, *S*al alone is expressed in the future black spots. CRISPR knockouts of *Dll* and *sal* showed that each gene is required for the development of melanic wing pattern elements, and repressing pteridine granule formation, in the areas where they are expressed. We conclude that both genes likely play ancestral roles in organising distal butterfly wing patterns, across pierid and nymphalid butterflies, but are unlikely to be differentiating signalling centers in pierids black spots. The genetic and developmental mechanisms that set up the location of spots and eyespots are likely distinct in each lineage.

## Background

Butterfly wings exhibit an astounding diversity of patterns shaped by their roles in thermoregulation (Kingsolver, 1985; Stoehr & Goux, 2008), mate choice (Silberglied & Taylor, 1978; Silberglied, 1984; Fordyce et al., 2002), and predator deterrence (Uésugi, 1996; Finkbeiner et al., 2014; De Bona et al., 2015). Of these wing patterns, eyespots, with their concentric rings of contrasting colors, are arguably one of the most well-studied patterns for their ecological functional roles in predator avoidance and in mate signaling (Robertson & Monteiro, 2005; Stevens, 2005; Stevens et al., 2007; Merilaita et al., 2011; Prudic et al., 2015; Ho et al., 2016; Chan et al., 2021). It is also interesting that simpler traits, such as spots in pierid and lycaenid butterflies (Fordyce et al., 2002; Stoehr et al., 2016), have also been implicated in mate signaling, but the developmental similarities and evolutionary relationship between spots and eyespots have remained unclear.

It is unclear whether nymphalid eyespots and pierid spots share similar origins. A study examining the phylogenetic distribution of spots and eyespots across the nymphalids, and a few outgroups suggested that eyespots replaced nymphalid spot patterns that were already present in specific wing sectors (Oliver et al., 2014). While we do not know whether both pierid and nymphalid spots share any degree of homology, it remains a possibility that the two may share similar developmental mechanisms. Alternatively, pierid spots may be homologous to submarginal bands of nymphalid butterflies as proposed by Schwanwitsch (Schwanwitsch, 1956) and Shapiro (Shapiro, 1984). In this proposal that is founded in comparative morphological work, pierid spots are not part of the *border ocelli* (eyespots) system but are rather positional homologs of more distal wing pattern elements (Fig 1). Schwanwitsch (Schwanwitsch, 1956) assigned the simpler spots of pierids as homologs to the Externa III (EIII), as did Nijhout (Nijhout, 1991), who classified these patterns as ‘*parafocal elements*’. Unfortunately, little is known about the developmental basis of spots, as well as other melanic wing patterns in pierids, for a proper evaluation of these two alternative hypotheses at a more mechanistic level.

**Figure 1.**
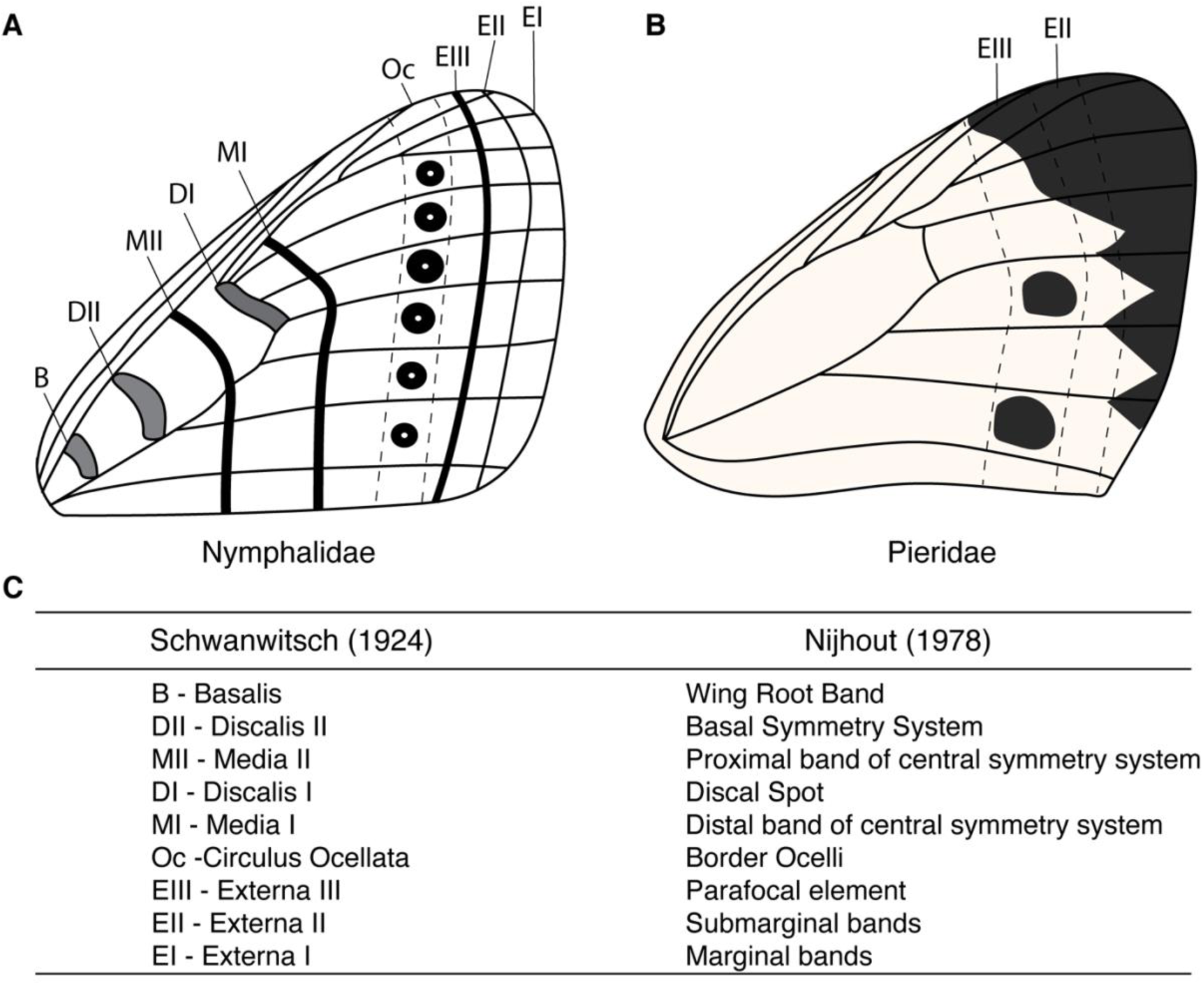
Schematic of wing patterns found on the wings of nymphalid and pierid butterflies. A) The nymphalid ground plan (NGP), a representation of the maximal number of pattern elements found in the wings of nymphalid butterflies, as devised by Schwanwitsch (Schwanwitsch, 1924). B) The NGP was subsequently extended and applied to the analyses of wing patterns of butterflies belonging to other families. Pierid butterflies were noted to have reduced wing patterns, with their wing spots thought to be positional homologs of the EIII band, also known as the parafocal element. C) Nomenclature of terms used in different versions of the NGP (Schwanwitsch, 1924; Nijhout, 1978).

The few experiments that have been performed in pierids indicate that spots show some differences but also some similarities to eyespots in terms of their development. Damage applied to the centre of eyespots and spots, in early pupal development, reduces the size of the respective patterns, suggesting that these cells might be important signalling cells in both cases (Nijhout, 1980; Stoehr et al., 2013). On the other hand, spots in pierids and eyespots in nymphalids show differences in the expression of a few candidate genes, as well as in cellular arrangements, at an earlier stage of development when those central cells should be differentiating. At the late larval stage, several genes required for eyespot center differentiation in nymphalids, including the transcription factors Distal-less (Dll) and Spalt (Sal) (Connahs et al., 2019; Murugesan et al., 2022), are absent from the presumptive spot centers of *Pieris rapae* butterflies (Monteiro et al., 2006; Saenko et al., 2011; Oliver et al., 2012). Furthermore, these two genes are hypothesized to be part of a reaction-diffusion mechanism that differentiates these central cells in nymphalids in each wing sector bordered by veins (Connahs et al., 2019). This group of cells, called the focus, are more densely packed and slightly raised from the wing plane relative to other epidermal cells (Iwasaki et al., 2017). In pierids, however, no such reaction-diffusion mechanism has been proposed for spot center differentiation, and the cells at the center of these spots resemble cells elsewhere on the wing. At early pupal stages of development, however, both Dll and Sal proteins are required for the differentiation of the black scales in eyespots of *B. anynana* (Connahs et al., 2019; Murugesan et al., 2022), and Sal protein, but not Dll, has also been associated with melanic scale patterns, including spots, in several pierids (Monteiro et al., 2006; Stoehr et al., 2013). However, the function of either gene has not been tested outside of nymphalids. In addition, to date, no studies have managed to functionally identify the up-stream signals that activate *Dll* and *sal* in melanic regions of either nymphalid eyespots or pierid spots.

Both *Dll* and *sal* have also been implicated in the development of melanic color patterns in other areas of nymphalid wings, and *sal* in the larval integument of papilionids. *Dll* is required for the background brown color in *B. anynana* wings (Connahs et al., 2019), and both genes are required for the development of pattern elements along the parafocal, marginal, and submarginal wing bands of numerous nymphalid species (Zhang & Reed, 2016; Connahs et al., 2019; Reed et al., 2020). Aside from wings, *sal* is also expressed in melanic regions of eyespot patterns on the larval epidermis of *Papilio xuthus* (Futahashi et al., 2012). This suggests that *sal*, and perhaps also *Dll*, may play a role in the development of melanic patterns outside nymphalids.

Here we test the function of both *Dll* and *sal* in pierid wing pattern development. We use CRISPR-Cas9 to target those genes in *Pieris canidia*, the Indian cabbage white. We also examine the expression of these transcription factors in a few additional nymphalid species that have spots, instead of eyespots, and explore the expression of Armadillo (Arm) protein and *decapentaplegic* mRNA, two possible up-stream activators of *Dll* and *sal* in both larvae and early pupae of *P. canidia*.

## Results

### Presence of Distal-less and Spalt proteins in *B. anynana* and *P. canidia*

We examined the distribution patterns of Dll proteins for both larval and 24 h pupal wings of *B. anynana* and *P. canidia* (Fig 2). Larval wing discs of both species showed strong levels of Dll along the wing margin, and in midline finger-like projections from the margin, between developing veins (Fig 2A & 2A’). Levels of Dll protein were higher in a cluster of cells at the end of these fingers in *B. anynana* larval and pupal wings but not in *P. canidia* (Fig 2A & Fig 2C). In *P. canidia* larval and pupal wings, Dll levels continue to be high in mid-line projections in individual wing sectors (Fig 2B’, 2D’). These findings are consistent with previous studies done in a closely related species, *Pieris rapae* (Reed & Serfas, 2004; Monteiro et al., 2006). A novel observation, however, is that Dll is also present in areas along the wing margin containing the black chevrons, and in the wing apex, mapping to the areas of melanized scales at these two locations (Fig 2I & 2I’).

**Figure 2.**
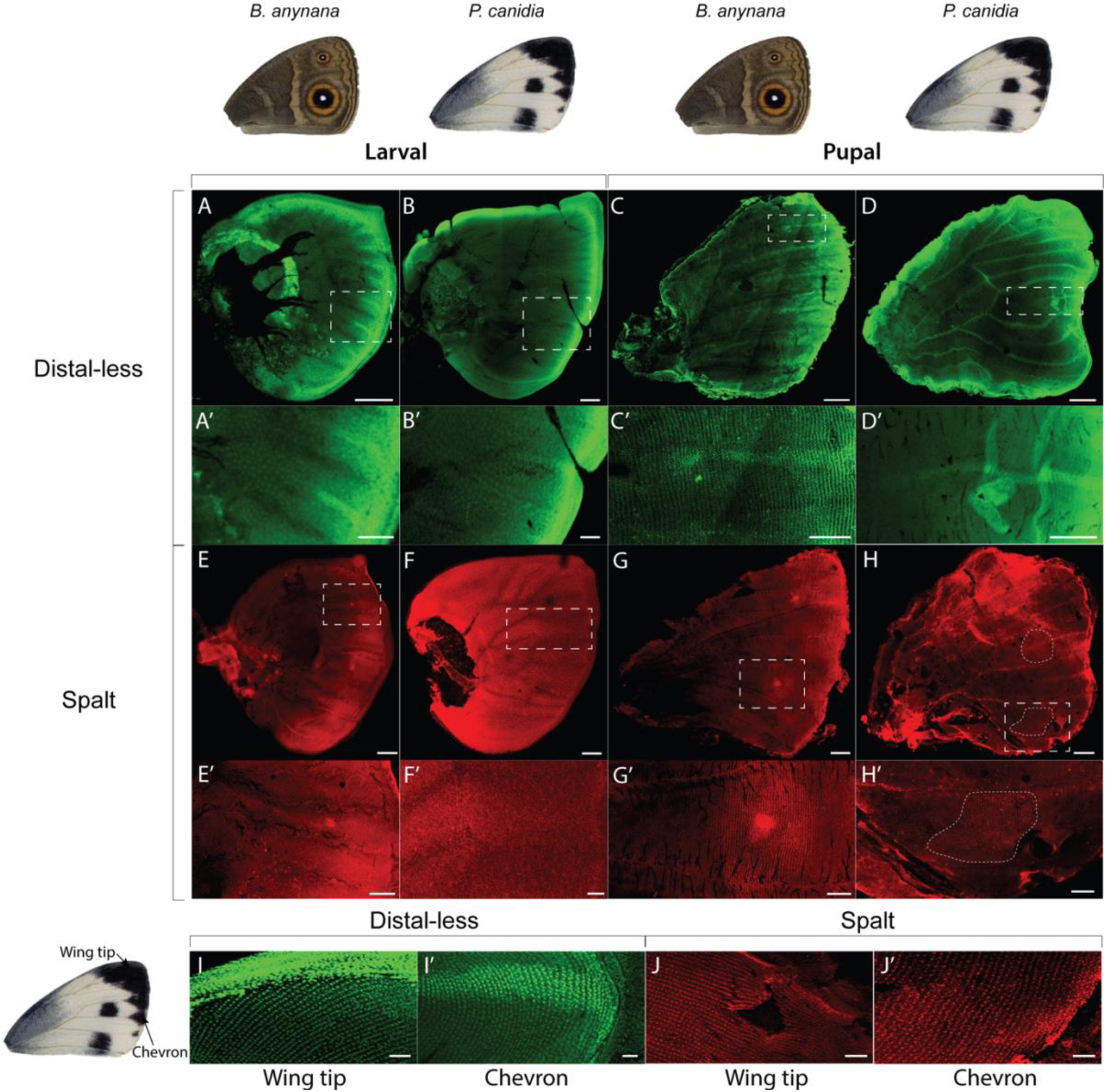
Immunostainings of Distal-less and Spalt proteins in larval and pupal wings. A-D’) Dll protein is present in late fifth instar larval and 24-26 h pupal wing discs. A, A’, C & C’) In *B. anynana* larval and pupal wings, Dll is observed between veins as finger-like projections from the wing margin, ending with a discrete focus at the proximal tip of the fingers, that corresponds to the eyespot centres. In pupal stages of development, Dll becomes additionally observed in cells that correspond to the black scales of the eyespot pattern. B, B’, D & D’) In *P. canidia*, intervein finger-like projections of Dll protein are observed but with no discrete foci at the tips of the fingers. E-H’). Sal protein is present in late fifth instar larval and 24-26 h pupal wings discs. E, E’, G, & G’) In *B. anynana*, Sal protein is observed in eyespot foci during the larval stage. Like Dll, Sal becomes additionally observed in the cells that map to the black scales in the eyespots during pupal wing development. F, F’, H, & H’) In *P. canidia*, there is no cluster of cells in the middle of the spot pattern that is expressing higher levels of Sal proteins in larval wings, and Sal is present in the cells that map to the black scales in spots in 24 h pupal wings. I, I’, J, J’) Dll and Sal proteins are also observed in cells that will become black scales located along the wing margin at both the wing tips and in the chevron patterns along the wing margin in *P. canidia*. Note the strong punctate nuclear staining of scale-building cells taken at 20x magnification. Scale bars for (C, D, G, and H – 500 μm); (A, B, B’, C’, D’, E, F, G’ and H’ – 200 μm); (E’ and F’ – 100 μm); (A’, I, I’, J & J’) – 50 μm)

The presence of Sal proteins was also examined for both species at the same time points in larval and pupal wings. In a similar manner to Dll, Sal proteins were present in the eyespot foci in late larval wings of *B. anynana* (Fig 2E, 2E’) but absent from spot centers in *P. canidia* (Fig, 2F, 2F’). In 24 h pupal wings, Sal was additionally observed in the scale-building cells that map to the black scales of an eyespot (Fig 2G’). In *P. canidia*, Sal was observed in the scale-building cells that map to all the densely melanised areas on the wing, including the black spots, the chevrons at the wing margin, and the apex of the wing (Fig 2H’, 2J & 2J’). However, spot centers did not have elevated levels of Sal, nor did these central cells appear distinct from surrounding spot cells, as they do in eyespots. These results are similar to those previously described for other pierids (Monteiro et al., 2006; Stoehr et al., 2013).

The protein localizations of Dll and Sal in three other nymphalid species were like those observed in *B. anynana*. Dll and Sal were present in the focal cells of future eyespots (of *Vindula dejone*) and spots (of *Hypolimnas bolina jacintha* and *Cethosia cyane*) and along the submarginal wing patterns during the larval stage (Fig 3). This pattern persisted in 24 h pupal wings but the two proteins were additionally present in a few surrounding scale-building cells that map to black pattern elements in an eyespot or spot. The simple white spots of *Hypolimnas bolina* are likely equivalent to central cells of an eyespot that have become reduced to a single ring/spot of color with just a few black cells around them.

**Figure 3.**
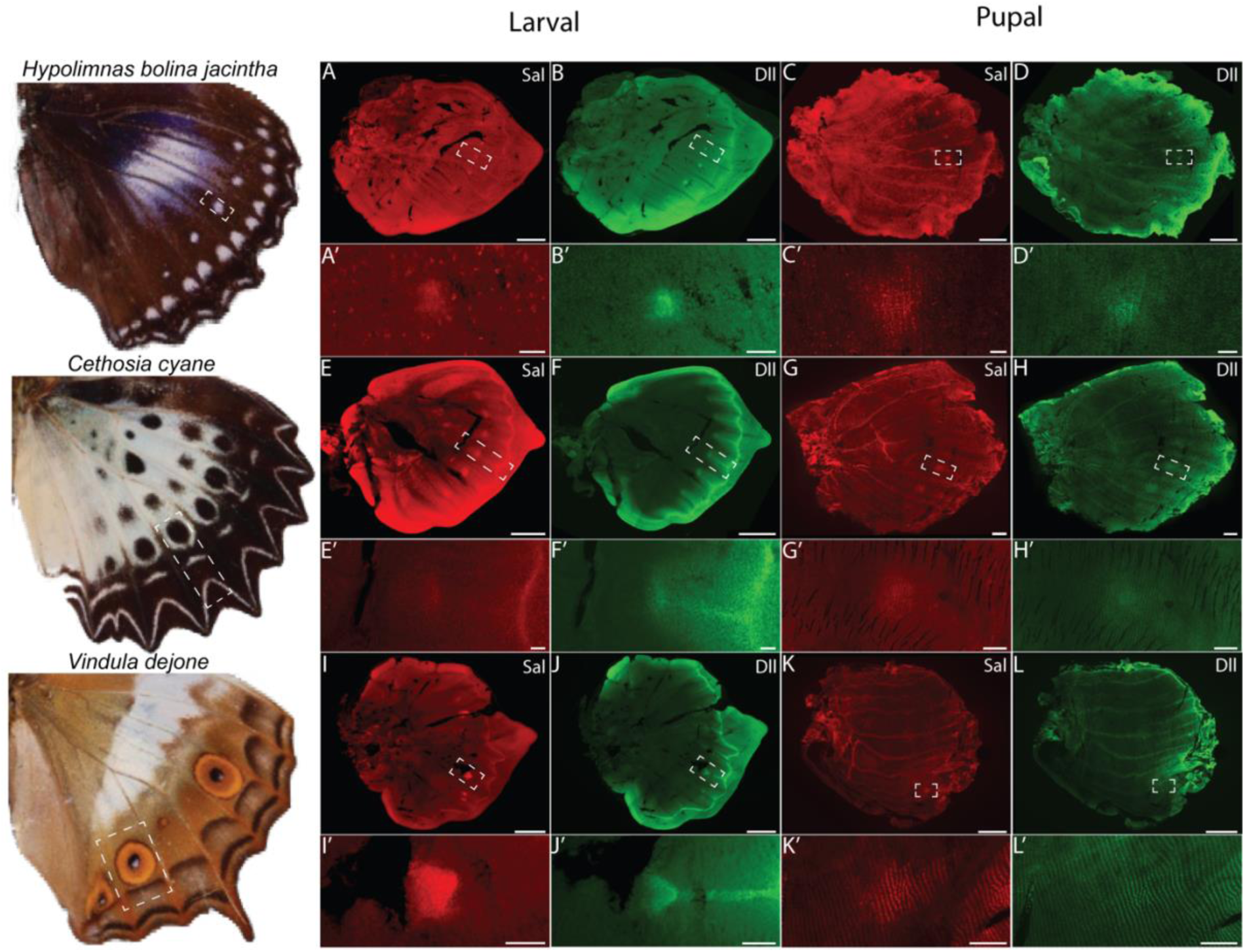
Immunostainings of Distal-less and Spalt proteins in other nymphalids with spot and eyespot patterns. In all species surveyed here, both Dll and Sal proteins are present in spots and eyespot patterns in late fifth instar larval and 24-28 h pupal wings. Note that both proteins are also expressed in wing patterns that map to parafocal, submarginal and marginal pattern systems as outlined in the nymphalid ground plan. Scale bars for (K and L – 1500 μm); (C and D – 1000 μm); (A, B, E, F, G, H, I, and J – 500 μm); (G’ and H’ – 200 μm); (A’, B’, C’, D’, E’, F’, I’, J’, K’ and L’ – 50 μm). The expression of Sal on the right in panel K’ corresponds to another, more posterior eyespot.

### Presence of Armadillo (Arm) and expression of *decapentaplegic* (*dpp*) in *B. anynana* and *P. canidia*

In the *Drosophila* wing margin, *Dll* is a downstream target of Wnt signalling (Campbell & Tomlinson, 1998), whereas in the center of the wing, *sal* is a target of Dpp signalling (Barrio & de Celis, 2004). To investigate whether Wnt and Dpp signalling could be upstream of the melanic patterns in *P. canidia*, we performed immunostainings targeting the protein Armadillo (Arm), a signal transducer of canonical Wnt signalling (Wodarz & Nusse, 1998), and performed *in situ* hybridizations with a probe against *dpp*. We found Arm present in the wing margin and in finger-like patterns from the wing margin in both *B. anynana* (as previously described in (Connahs et al., 2019)) and *P. canidia* (Fig 4A & 4B). However, Arm was present in the eyespot centers in *B. anynana* but not in spot-like patterns in *P. canidia* during both larval and pupal stages (Fig 3A’, 4B’, 4C’ & 4D’). This suggests that Wnt signaling is stable and active in *B. anynana* eyespot centers but not in *P. canidia* spot centers. In *B. anynana, dpp* is present in cells flanking the veins and along the anterior-posterior (AP) boundary (as previously described in (Connahs et al., 2019; Banerjee & Monteiro, 2020b), and later in eyespot centers in 18 h pupal wings (Fig 4E & 4G). In *P. canidia* larval wings, *dpp* is expressed strongly along the veins and the border lacuna, parallel to the wing margin. No *dpp* was detected in areas mapping to the spot pattern in 18 h pupal wings (Fig 4F & 4H).

**Figure 4.**
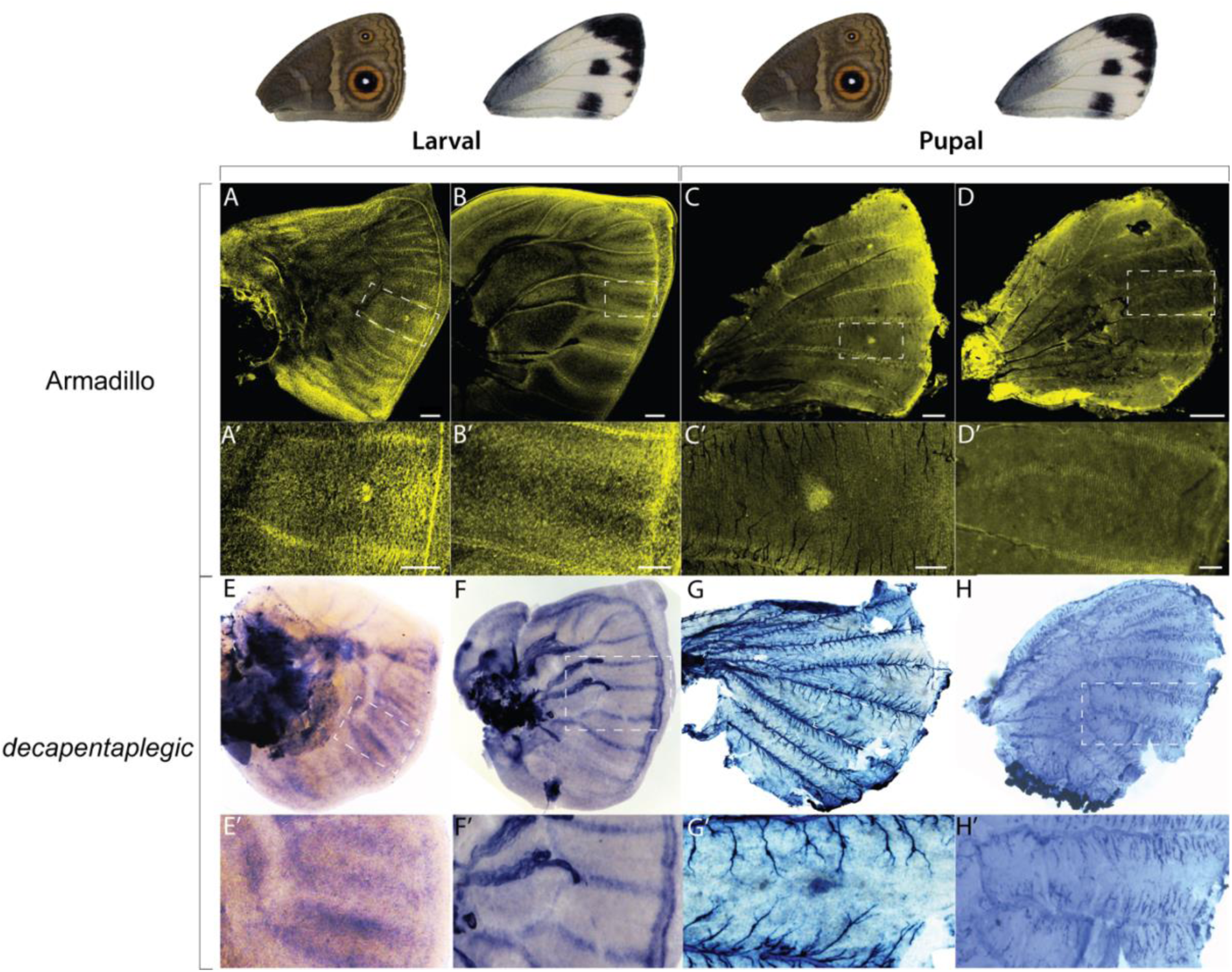
Expression of Armadillo (Arm) protein, and *decapentaplegic* (*dpp*) mRNA in larval and pupal wings. A, A’, B, B’, C, C’, D & D’) Distribution of Arm protein in late fifth instar larval and 20 h pupal wings. A, A’, C & C’) In *B. anynana*, Arm is present along the wing margin and in eyespot foci in both larval and pupal wings. B & B’) In *P. canidia* larval wings, Arm is present between veins in finger-like projections, in a similar pattern to that of Distal-less. D & D’) Arm is not present in the black spots of *P. canidia* in 20 h pupal wings. E, E’, F, F’, G, G’, H & H’) Localisation of *dpp mRNA* transcripts in late fifth instar larval, 18 h pupal wings (*B. anynana*) and 18 h pupal wings (*P. canidia*). E, E’, G, & G’) *dpp* is expressed in areas flanking the veins in *B. anynana* larval wing discs and is absent from eyespot foci at this stage. *dpp* is expressed in eyespot foci in 18 h pupal wings. F & F’’) *dpp* is expressed strongly along veins and along the border lacuna in *P. canidia* larval wings. H & H’) *dpp* is not expressed in the center of spot patterns in 18 h pierid pupal wings. The wing used for *dpp in-situ* hybridisation in Fig 1G and 1G’ is a *B. anynana* hindwing.

### Both Dll and Sal regulate melanic wing patterns in *P. canidia*

To test the function of *Dll* in spot development and melanisation, we targeted both exons 2 and 3 using the CRISPR/Cas9 system (Fig 5A). Consistent with the immunostaining results for Dll, melanic wing patterns located along the wing tip and in chevrons along the wing margin were disrupted (Fig 5C). We did not observe any disruptions to the black spot pattern, at least within the small number of *Dll* mutants that were obtained in this study. In the affected areas, black scales were transformed into white scales. In two of the crispants, however, both ground and cover scales were missing from the affected regions (Fig 5D & 5F).

**Figure 5.**
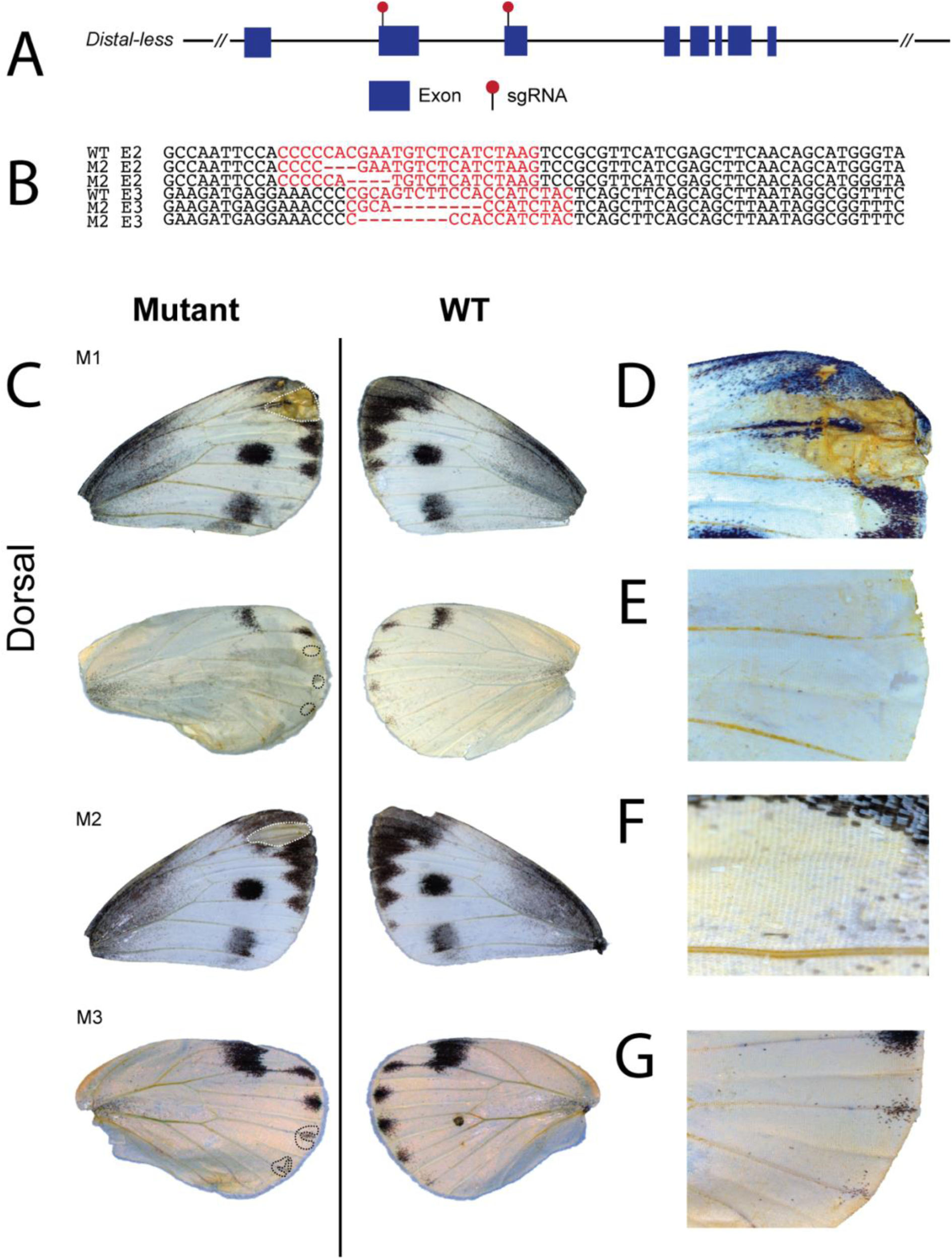
*Distal-less* functions in the development of wing margin melanic scale development in *P. canidia*. A) Structure of the *Distal-less* locus and location of the two sgRNAs used to disrupt the locus in exons 2 (E2) and exon 3 (E3) (red pins). B) *Dll* crispants had indels in both E2 and E3 that were detected using Next-Generation sequencing. C) Various *Dll* crispants generated through CRISPR/Cas9 of both E2 and E3. Phenotypes include disrupted scale development and possible loss of melanism as supported by aberrant phenotypes obtained in D) defective wing margin with loss of both black and white scales within the affected area, F) loss of black and white scales in the wing apex, and E & G) transformation of black scales in chevron areas to white scales. (D-G) Close-up of the mosaic area affected by the CRISPR knock-out experiments. Crispants shown here were affected by disruptions in both Exons 2 and 3.

To test the function of *sal* in spot development and in scale melanisation, we targeted exon 2 with the CRISPR/Cas9 system. The resulting mosaic phenotypes support a role for *sal* in scale melanisation in the spots and chevrons along the wing margin. We observed missing spots on both dorsal and ventral surfaces of forewings, fragmented spots, and a missing black wing marginal chevron in a single individual (Fig 6C, M8). Black scales in these areas were transformed into white scales. In addition, we saw one individual with less melanised scales (Fig 6C, M9).

**Figure 6.**
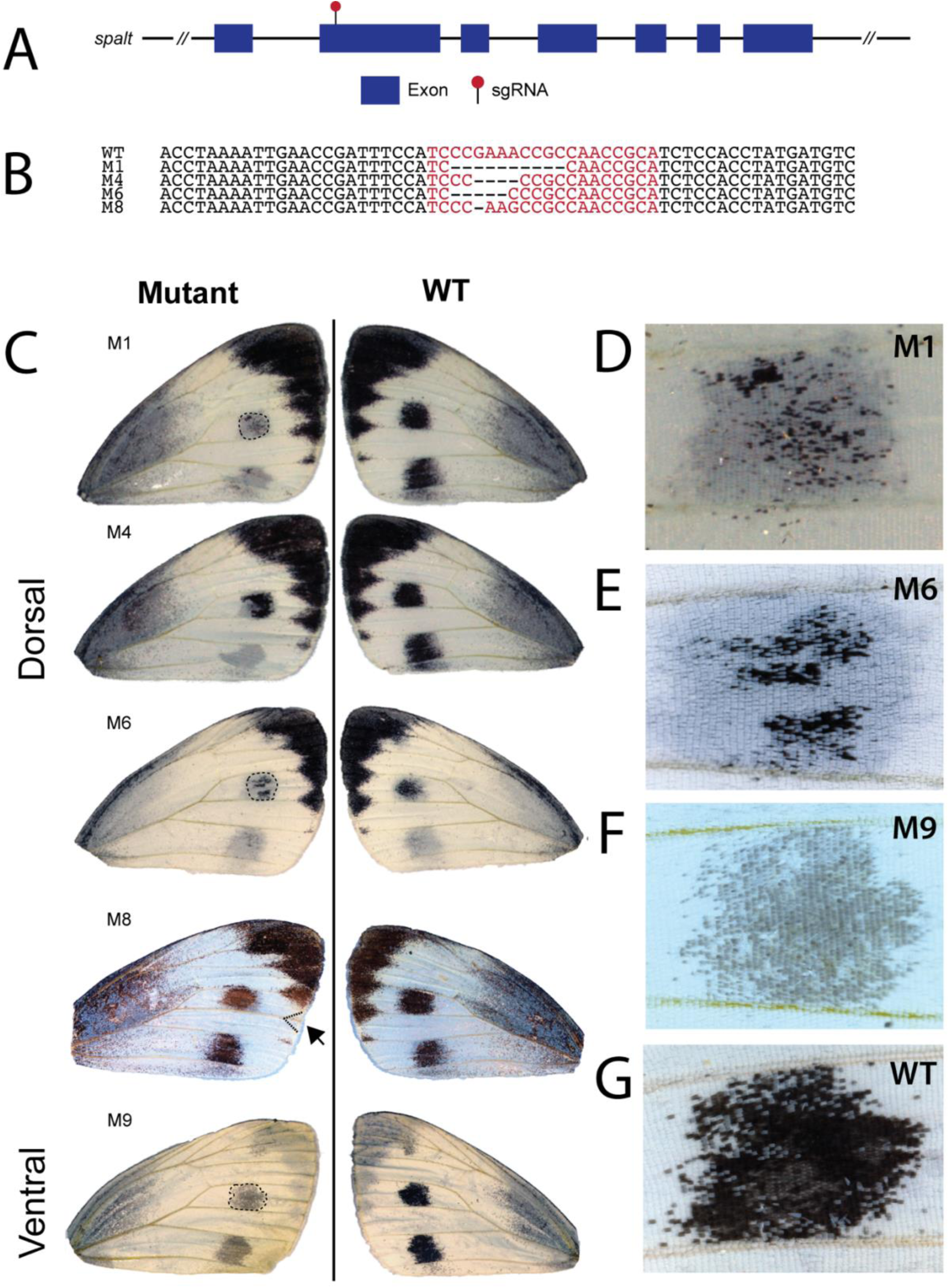
*spalt* functions in black scale development in *P. canidia*. A) Structure of the *spalt* locus and area targeted by the sgRNA (red pin). B) *spalt* crispants had indels in the target region that were detected using Sanger sequencing. C) Various *spalt* crispants (mosaic mutants) generated through CRISPR/Cas9. Phenotypes include missing spots or missing black scales in spots, disrupted Cu2 veins, missing black chevrons located along the wing margin (M8), and less melanised spots (M9). D-F) Close up of mosaic areas affected. G) Close up of black spot pattern in wild-type *P. canidia*.

Individual scales of *Dll* and *sal* mutants and wild-type butterflies were then closely examined using Scanning Electron Microscopy (SEM) to look for any changes in scale structure that might be under the regulation of either gene. Wild-type black scales had little to no pigment granules present, in contrast to white scales (Fig 7A). In both *Dll* and *sal* mutants, black scales that transformed into white scales contained dense rows of ovoid-like pigment granules deposited along the cross-ribs (Fig 7B & 7C), resembling WT white scales. The scales of the *spalt* crispant that displayed less melanised scales in the black spot region (Fig 7D) were intermediate in colour and in morphology – the windows were not completely open, and remnants of upper lamina were observed along the crossribs as compared to Wt black scales (Fig. 7D). Pigment granules were also scattered within the scale lumen.

**Figure 7.**
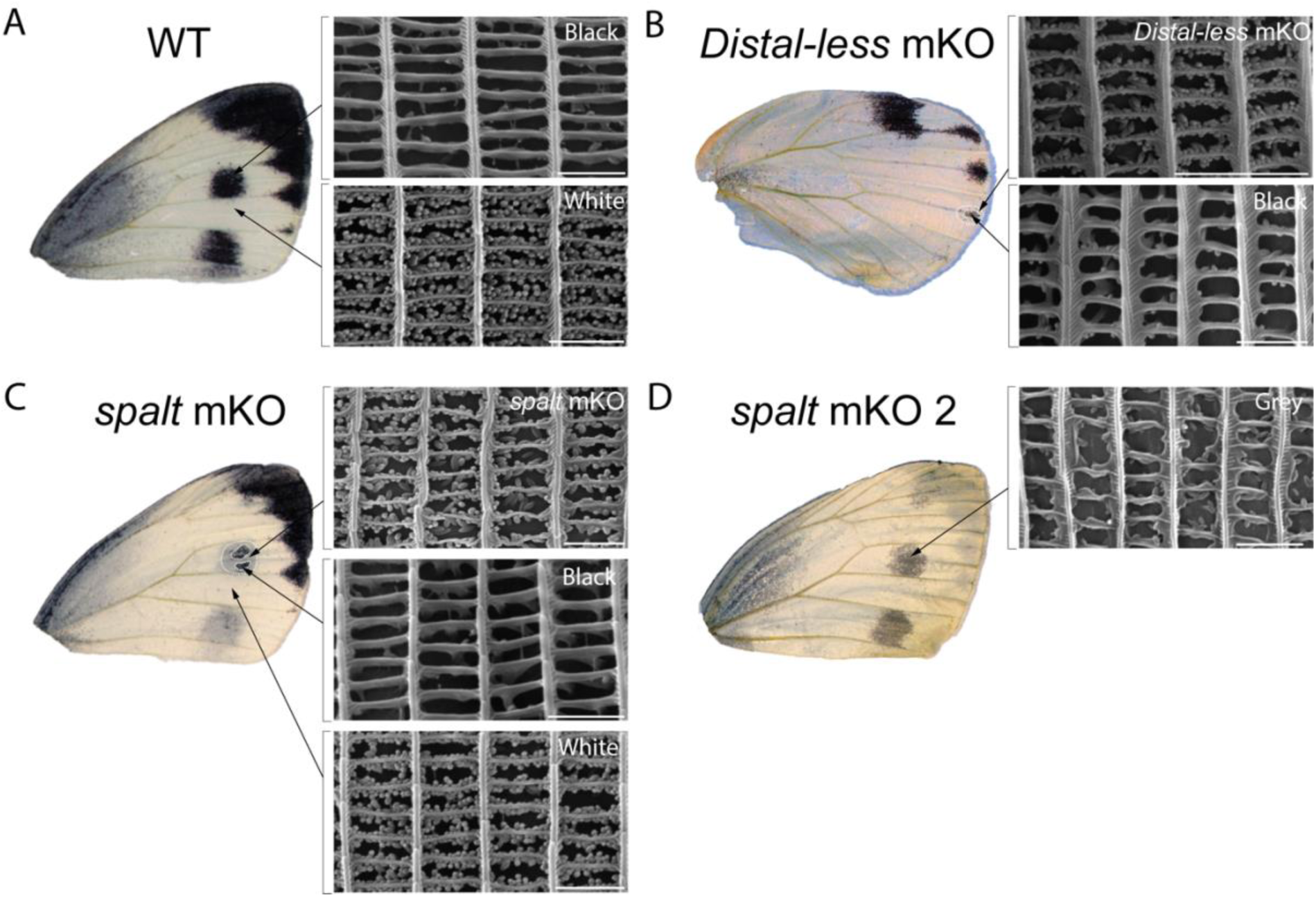
Melanized scales that become white scales acquire pterin pigment granules visible under scanning electron microscopy. Individual *P. canidia* scales were removed from wild-type black and white regions, as well as from *spalt* mKO, and *Distal-less* mKO affected regions. A) SEM images of a black scale and a white scale removed from the forewing of wild-type *P. canidia*. Close-up of a black scale showing no pigment granules present along the cross-ribs of the scale. Pigment granules are present in great numbers in white scales. B) SEM images of black scales and white scales removed from a *Dll* crispant. This crispant had greatly reduced spots on its hindwing. Scales that lost melanin pigments showed a morphology resembling that of WT white scales. C) SEM images of black and white scales removed from a *spalt* crispant. *The* SEM image labelled as *spalt mKO* showed a close-up view of a scale (originally black) removed from the CRISPR/Cas9 mosaic knockout area. Black scales converted into white scales with pigment granules, resembling those of wild-type white scales. D) SEM images of a *spalt* mutant that displayed an intermediate scale phenotype with less melanised scales in the black spot region. The morphology of these grey scales resembles that of WT black scales, but windows of these scales were not fully opened and there remains residues of the upper lamina. Scale bars: 2 μm.

## Discussion

The extent of wing pattern homologies shared between different butterfly families remains elusive due to a lack of functional genetic studies outside of the nymphalids. Here, we provide functional evidence for a deeply conserved role of two transcription factors, *Distal-less* and *spalt*, as pattern organisers of distal butterfly wing patterns. We also show that *spalt* behaves like a ‘switch gene’ for pierid wing patterns, mediating eventual scale colour fates between pterins and melanin, much like a previously reported function for the gene *optix* (Zhang et al., 2017). Lastly, we lend further support to the hypothesis that pierid spots are unlikely to be positional homologs of nymphalid eyespots. Unlike eyespot center differentiation, spot differentiation does not depend on the expression of either *Dll* or *sal* at the center of the pattern during the larval stages of development.

Previous research suggested that eyespots may have derived from pre-existing nymphalid spot patterns (Oliver et al., 2014), but genes previously associated with nymphalid eyespot patterns were not found in spot patterns of other butterfly families, apart from *sal* (Monteiro et al., 2006; Bhardwaj et al., 2020). Here we show that both *Dll* and *sal* have deeply conserved roles in organising distal wing pattern elements in lepidopteran wings, predating the divergence of nymphalid and pierid butterflies. *sal* knockouts showed disrupted black spots and marginal markings, whereas *Dll* knockouts affected both scale development as well as melanic patterns located along the wing tip and wing margins of both forewings and hindwings.

While both genes are required for the formation of black marginal chevrons and wing tips, *sal* alone is sufficient for the development of wing spots in *P. canidia*. We postulate that *Dll* is likely working upstream of *sal* in areas where the two genes are co-expressed, but not in the black spot area of *P. canidia*. The regulatory interaction between *sal* and *Dll* has been inferred from mutants and from functional work in *B. anynana*. In wildtype *B. anynana*, both *spalt* and *Dll* are co-expressed in the white centers, in the chevron patterns, and in the black scales of an eyespot during the pupal stages (Brunetti et al., 2001; Murugesan et al., 2022). In the larval stages, *Dll* is required for *sal* activation in the eyespot centers and marginal chevrons, whereas *sal* is not required to regulate *Dll* (Murugesan et al., 2022). In the pupal stages, *Dll* is required for melanin pigment production in the black scales and in background brown wing scales (Connahs et al., 2019), whereas *sal* is required to repress *optix* from becoming expressed in the central black disc of an eyespot, and from turning these scales into orange scales (Banerjee et al., 2021). Further, in *Goldeneye B. anynana* mutants, which had its black scales replaced by orange scales within the eyespot pattern, Dll proteins persisted while Sal proteins were absent (Brunetti et al., 2001; Murugesan et al., 2022). This suggests that *Dll* is either working upstream of *sal*, in both larval and pupal stages, or parallel to *sal* in the pupal stage in *B. anynana*. In this species, both *Dll and sal* are required for the development of black scales in eyespots. This same circuit might also be deployed in the tips and black chevrons of *P. canidia* pupal wings, but additional work will be necessary to confirm this.

It is plausible that in the case of pierid spots, both genes may be directly or indirectly regulating enzymes from the melanin biosynthesis pathway. If so, the developmental mechanism underlying the differentiation of melanic spots and melanic areas in eyespots may be homologous in this context, with the same genes performing a similar function i.e., differentiating black scales in both traits. We still do not know how melanin pathway genes are being regulated by either *Dll* or *sal* nor do we know the upstream signal(s) that both genes are responding to in lepidopterans. Previous studies have shown that expression of both Dll and Sal proteins also correlate with patterns of different colour states on the wing. In 16-24h pupal wings, expression of Sal protein spatially maps to pale-colored non-eyespot marginal wing patterns of nymphalids (Reed et al., 2020) while both Dll and Sal proteins are expressed in silver scales along the wing margin in the lycaenid butterfly, *Lycaeides melissa* (Brunetti et al., 2001). Thus, both *Dll* and *sal* may be ancestral pattern organisers working within the distal part of the wing, operating independently of melanic fate. Nevertheless, future studies should try to unravel the possible regulatory connections between *Dll* and *sal* and downstream melanin biosynthesis genes, including investigating whether intermediate transcription factors mediate this link.

Similar to a previously reported gene, *optix* (Zhang et al., 2017), *spalt* may be functioning as a ‘switch’ gene that represses the pterin biosynthesis pathway (white) while activating the melanin biosynthesis pathway (black). If *spalt* was purely an upstream activator of genes involved in melanin synthesis, we would expect to see scale morphology of mutant scales resembling those of the flanking black scales that were unaffected by the CRISPR/Cas9 knockout. However, when *spalt* mutant scales were examined using SEM, we observed numerous pigment granules densely arranged along the cross-ribs, closely resembling the structures found in wildtype white scales. White scales of pierid butterflies differ from those of other butterfly species in that many ovoid beads are attached to the cross-ribs of each scale (Ghiradella et al., 1972; Stavenga et al., 2004; Wilts et al., 2017). These beads contain leucopterin, a class of heterocyclic pigment that absorbs exclusively in the ultraviolet range. When coupled with the strong light-scattering properties of these beads, leucopterin filled granules cause scales to appear white (Wilts et al., 2011). Our examination of the poorly melanised spot that was likely derived from a hypomorphic allele of *sal*, or perhaps a heterozygote crispant clone, suggests that intermediate scale colors (grey) and morphologies are possible (Fig 7D). This mutant suggests that intermediate levels of Sal protein might be insufficient for complete downregulation of the pteridine pathway and for complete up-regulation of the melanin pathway.

Dll mutant clones displayed two phenotypes, loss of all scales and a change in scale color from black to white along marginal pattern elements. The loss of both cover and ground scales, lends further support to butterfly scales being a derived form of a sensory bristle (Galant et al., 1998) that requires *Dll* for its development (Panganiban, 2000). This corroborates a previous finding by (Connahs et al., 2019) whereby loss of scales was also observed in *Dll* crispants in *B. anynana*. The transformation of black to white scales may be connected to hypomorphic alleles of *Dll*, or perhaps to heterozygote crispant clones. It is tempting to speculate that like *sal, Dll* might also regulate two different pigment pathways simultaneously. However, it is more likely *Dll* was working upstream of *sal* in the wing marginal patterns and that knocking out *Dll* resulted in the downregulation of *sal*, leading to the formation of ectopic pigment granules. This is also supported by the observation that knockouts of *sal* alone, in spots, produces the scale color switch phenotype.

Nymphalid eyespot evolution, however, may have relied on the novel larval expression of *Dll* and *sal* in the foci at the tips of intervein fingers, after the divergence of nymphalids from pierids. This novel expression may have taken place through a gradual increase of *Dll* expression that can promote a stable expression of *Dll* at the foci via a reaction-diffusion mechanism (Connahs et al., 2019) (Fig 8). Higher Dll levels, in turn, may be dependent on Wnt and dpp signals which become anti-colocalized at late stages of eyespot focus differentiation, again via the same reaction-diffusion process (Connahs et al., 2019) (Fig 8). In *P. canidia*, Armadillo protein patterns were quite similar to those observed in *B. anynana* but again, no Arm foci were detected at the end of the intervein fingers (Fig 4B’). The *dpp* pattern was also different in *P. canidia* and was not anti-colocalized with the Arm pattern (Fig 4F’). This suggests that a reaction-diffusion mechanism like that proposed for *B. anynana* is not taking place in *P. canidia* during mid-larval development.

**Figure 8.**
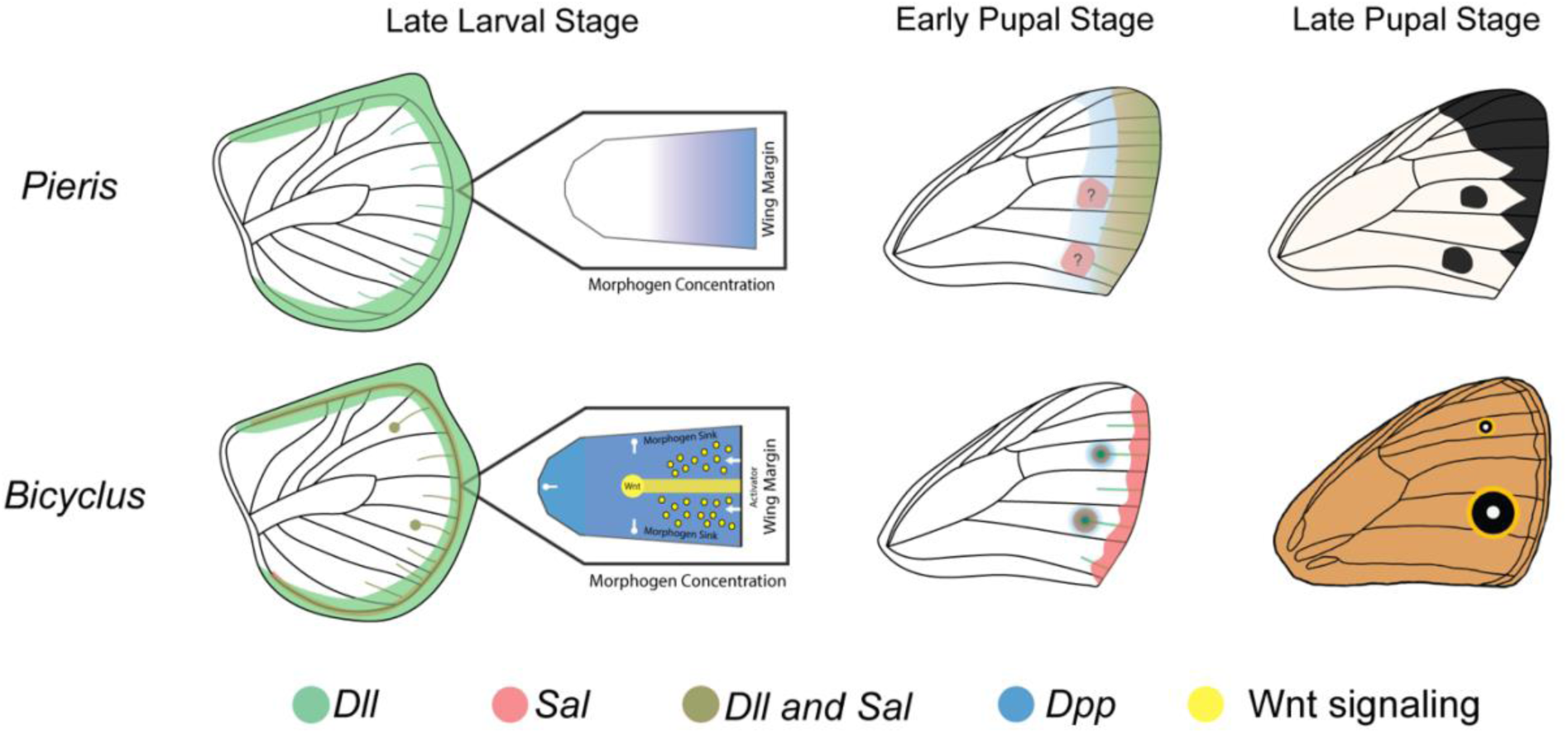
Possible roles of *Distal-less* and *spalt* in pierid spot and nymphalid eyespot development. In late larval wing discs of *B. anynana*, both *Dll* (green) and *sal* (orange) are co-expressed at high levels in the center of eyespots (Reed et al., 2020). However, in late larval wing discs of *P. canidia, Dll* and *sal* are not expressed in spot centers. Both *Dll* and *sal* are expressed in mid-line fingers encroaching inwards from the wing margin. Eyespot centers in *B. anynana* are likely established through a reaction diffusion mechanism involving Wnt and BMP signaling (Connahs et al., 2019). The absence of Arm proteins and *dpp* expression in *P. canidia* spot centers suggests that spots may not develop through the same mechanism. In nymphalid eyespots, *Dll* and *sal* respond to signals emanating from the foci. However, in early pupal stages, both Arm and *dpp* are absent in spot centers in pierids. There may be central signaling cells that are present in spot patterns that are activating downstream genes (i.e *sal*), but these central cells do not express Dll and Sal. An alternative model would be that *sal* is responding to a gradient of BMP ligands at specific thresholds (blue band in the early pupal stage). Inhibitory molecules (brown) secreted from the wing margin, as well as others expressed in specific wing sectors (not shown), would lead to *sal* expression and black spot markings of *P. canidia* in only specific wing sectors. In both butterfly lineages, *sal* likely plays an ancestral role in organising distal wing patterns as expression of Sal proteins have been observed along the marginal wing bands of *B. anynana* and *J. coenia* in early pupal wings (Reed et al., 2020).

The mechanism that sets up spots and black discs of color around eyespots, during the pupal stage, may also be distinct. During early pupal stages, no discernible Arm or *dpp* signals were observed in spot centers (Fig 4D’ & 4H’) as they were in eyespot centers (Fig. 4C’, 4G’). It is possible that *sal* in *P. canidia* may be responding to a gradient of BMP ligands such as *dpp* that is emanating from the wing margin. High levels of *dpp* expression were present along the wing margin of *P. canidia* larval wings (Fig 4F’), but not in *B. anynana* (Fig 4E’). Thus, we speculate that the role of *Dll* and *sal* in establishing nymphalid eyespot foci is novel and derived as compared to pierid spot development.

This derived role of *Dll* and *sal* as eyespot center organisers is supported by the fact that in late larval wings, the expression of both *Dll* and *sal* in the presumptive eyespot centers in nymphalid species is essential for eyespot development (Zhang & Reed, 2016; Connahs et al., 2019; Murugesan et al., 2022). Knockouts of *Dll* and *sal* in *B. anynana* that affected cells located in the eyespot center always led to the complete disappearance of an eyespot (Connahs et al., 2019; Murugesan et al., 2022). The expression of both genes, however, is absent from spot centers in pierid species during the larval stage (Reed & Serfas, 2004; Stoehr et al., 2013). Correspondingly, when scale cells located in the spot center were affected in *P. canidia spalt* knockout mutants, we did not observe entire spots disappearing. Instead, scattered areas of the spot retained melanised scales (Fig 6C).

Collectively, our results suggest that pierid spots are unlikely homologs of patterns in the *‘border ocelli’* band but may be positional homologs of more distal pattern elements with respect to nymphalid eyespots located within the ‘*EIII’* or ‘*parafocal elements’* banding systems. *Dll* and *sal* knockout mutants in nymphalid butterflies showed a disruption to both submarginal and marginal pattern elements (EI-III) (Zhang & Reed, 2016; Connahs et al., 2019; Reed et al., 2020). Given the classification of pierid spots as part of the EIII band, we expected that knocking out *Dll* in *P. canidia* should also result in disruption or missing spot patterns. However, we only observed disruptions along the black chevrons and wing tips, which are elements that correspond to the EI and EII bands. We speculate that *Dll* may not have a role in elaborating the EIII submarginal band in pierid wings, and that its function in organising the EIII band in nymphalids, may be a derived one, but comparative work will need to be done to validate this hypothesis.

The developmental mechanism of pierid spot differentiation is not yet fully understood. Pierid spots, like nymphalid eyespots, may rely on differentiated cells at their center to signal to surrounding cells to differentiate the complete spot pattern, as previously proposed (Stoehr et al., 2013). Alternatively, spots may be fragments of an anterior-posterior banding system that relies instead on activator signals spreading from the wing margin (Monteiro et al., 2006). More recent revisions of the NGP placed both eyespots and parafocal elements as part of the *Border Symmetry System* and heat shock experiments involving nymphalid species showed a fusion of these pattern elements (Otaki, 2012; Nijhout, 2017; Otaki, 2021). Both pattern elements may possibly arise from a common developmental origin. Regardless of the exact mechanism of spot development, our current experiments show that spots do not rely on *Dll* and *sal* being expressed at their center during the larval stages to differentiate.

## Conclusion

In this study, we tested the function of two transcription factors essential for nymphalid eyespot development, *Dll* and *sal*, in a basal butterfly lineage with primitive spots and other melanic patterns on its wings, *P. canidia*. Our work suggests that each transcription factor is required for the differentiation of distinct melanic elements in this species, including the spots, but these genes have no role in positioning spots on the wing. The mechanism of setting up the position spots and eyespots is likely to be distinct in the two lineages. Future work involving functional knockouts of other candidate genes or studying the expression profiles of some of these genes at additional time points will be able to shed additional light on the evolution of lepidopteran spot patterns.

## Materials and methods

### Animals

*Pieris canidia* used in this study were the descendants of wild-caught individuals from Singapore. Larvae were fed on potted *Brassica chinensis var*. parachinensis plants and adults on 10% sucrose solution. *Bicyclus anynana* larvae were fed on potted corn and adults on mashed banana. Both species were reared at 27°C and at 60% humidity under a 12:12 h light/dark photoperiod. All other species of butterflies used for comparative immunostainings work were reared at Entopia, a butterfly farm (Penang, Malaysia) under outdoor conditions.

### Immunostainings

Immunostainings were performed on 5^th^ instar larval wings and 16-30 h pupal wings dissected based on a protocol previously described by (Banerjee & Monteiro, 2020a) in 1X PBS at room temperature. Wings were fixed with 4% formaldehyde for 30 mins, washed with 1X PBS for four times at 10 mins, and transferred to 2 mL tubes filled with block buffer for blocking at 4°C for up to several months to reduce non-specific binding of the antibodies. Wing discs were then incubated in primary antibodies against Distal-less (1:200, mouse, a gift from Grace Boekhoff-Falk), and Spalt (1:10000, guinea-pig Sal GP66.1) overnight at 4°C, washed with multiple rounds of wash buffer, and stained in secondary antibodies anti-mouse AF488 (Invitrogen, #A28175) and anti-guinea pig AF555 (Invitrogen, **#**A-21435) at a concentration of 1:500. Stained wings were then washed with multiple rounds of wash buffer, away from light, and mounted on glass slides with an in-house mounting media. Images of the wings were taken with an Olympus FV3000 Confocal Laser Scanning Microscope. All buffer compositions are summarised in Table S2.

### Whole-mount *in-situ* hybridisation

*In-situ* hybridisations were performed on early to late 5^th^ instar larval wings and 16-18 h pupal wings dissected in 1X PBS at room temperature to prevent the crumpling of wings. The wings were fixed with 4% formaldehyde in PBST for 30 mins, digested with 1.25 μL of Proteinase-K in 1mL of 1X PBST for 5 mins on ice. The digestion reaction was stopped with a 2 mg/mL glycine solution in 1X PBST and followed with 3 washes of 1X PBST. Larval wings were removed from ice briefly for 5 mins and placed right back on ice to induce ‘puffing’ of the peripodial membrane for easier removal of the membrane using fine tip forceps. After removing the peripodial membrane, the wings were transferred to increasing concentrations of pre-hybridisation buffer in 1X PBST and incubated at 60°C for at least 1 h in pre-hybridisation buffer. Incubated wings were hybridised at 60°C with the probe (100 ng/μL) in a hybridisation buffer for 16-24 h. The next day, after incubation with the riboprobe, wings were washed with pre-hybridisation buffer for 5 × 10 mins at 60°C. The wings were then brought back to room temperature and transferred to 1X PBST gradually. 1X PBST was used to wash the wings for 2 × 5 mins, and wings were subsequently transferred for blocking for 1 h. Anti-digoxygenin was diluted in block buffer at a ratio of 1:3000 for incubation with the wings for 1 h. Once completed, the wings were washed with block buffer for 5 × 5 mins on a rotary shaker and transferred to an alkaline phosphatase buffer containing NBT-BCIP. Wings were left to incubate in the dark to develop color signal to the required intensity. A Leica DMS1000 microscope was used to image the stained wings. All buffer compositions are summarised in Table S3.

### CRISPR-Cas9

Knock-outs of the genes *Dll* and *sal* in *P. canidia*, were generated using the methods outlined in a previously published protocol (Banerjee & Monteiro, 2018). Single guide RNAs (sgRNAs) targeting the genomic regions of exons 2 and 3 of *Dll* and exon 2 of *sal* were designed using the webtool CHOPCHOP (Labun et al., 2019). For the gene *sal*, a total of 575 embryos were injected with a mixture containing 300 ng/μL of sgRNA (one guide) and 600 ng/μL of Cas9 protein (NEB, M0641) while for *Dll*, 357 embryos were injected with a mixture containing 100 ng/μL of sgRNAs (2 guides) and 300 ng/μL of Cas9 protein (Table S3).

Wild-type *P. canidia* laid eggs on a piece of parafilm that was wrapped around a small container that had its top covered with a piece of fresh cabbage leaf. The container was placed within the butterfly cage for up to 6 hours at a time to maximise the number of eggs collected. The parafilm and leaf were then removed from the container and transferred to a petri-dish for injection with the Cas9 injection mixture. Pieces of moist cotton wool were placed in each petri-dish post injection to avoid desiccation of injected eggs. Hatchlings were then directly transferred to *Brassica sp* plants and reared to adult eclosion. Upon emergence, the butterflies were frozen immediately in separate glassine envelopes and examined under the microscope for asymmetrical (left-right wing) phenotypic defects. Genomic DNA was isolated from the affected mosaic areas from CRISPR mutants, and indels were identified through Sanger and NGS sequencing.

### Scanning electron microscopy (SEM) imaging

Adult wing scales located in areas affected by the CRISPR experiment were individually picked with a needle and placed on carbon tape. All samples were sputter-coated with gold to increase conductivity and to reduce static surface charge. Samples were imaged using a JEOL JSM 6010LV Scanning Electron Microscope at 15-20 kV.

## Authors’ contributions

Both JW and AM conceptualise and designed the study. JW, TDB, AP, and KSS performed the experiments. JW analysed the data, and wrote the manuscript with input from AM. All authors read and approved the final manuscript.

## Acknowledgments

We thank Mr. BT Chin, Ms. Kuennie Lee, and Mr. Gan Gim Chuah (Entopia, Penang, Malaysia) for their support and for supplying some of the butterflies used in these experiments and Christopher Wheat (Stockholm University, Sweden) for providing the initial genomic sequences of *Dll* and *sal* in *Pieris canidia*. We also thank Ms. Tong Yan (Centre for BioImaging Sciences, National University of Singapore) for providing us help in the acquisition of the immunostaining images, Sree Vaishnavi and Gianluca Grenci (MBI, National University of Singapore) for access and help with SEM.

## Supplementary Files

**Supplemental Table 1.**
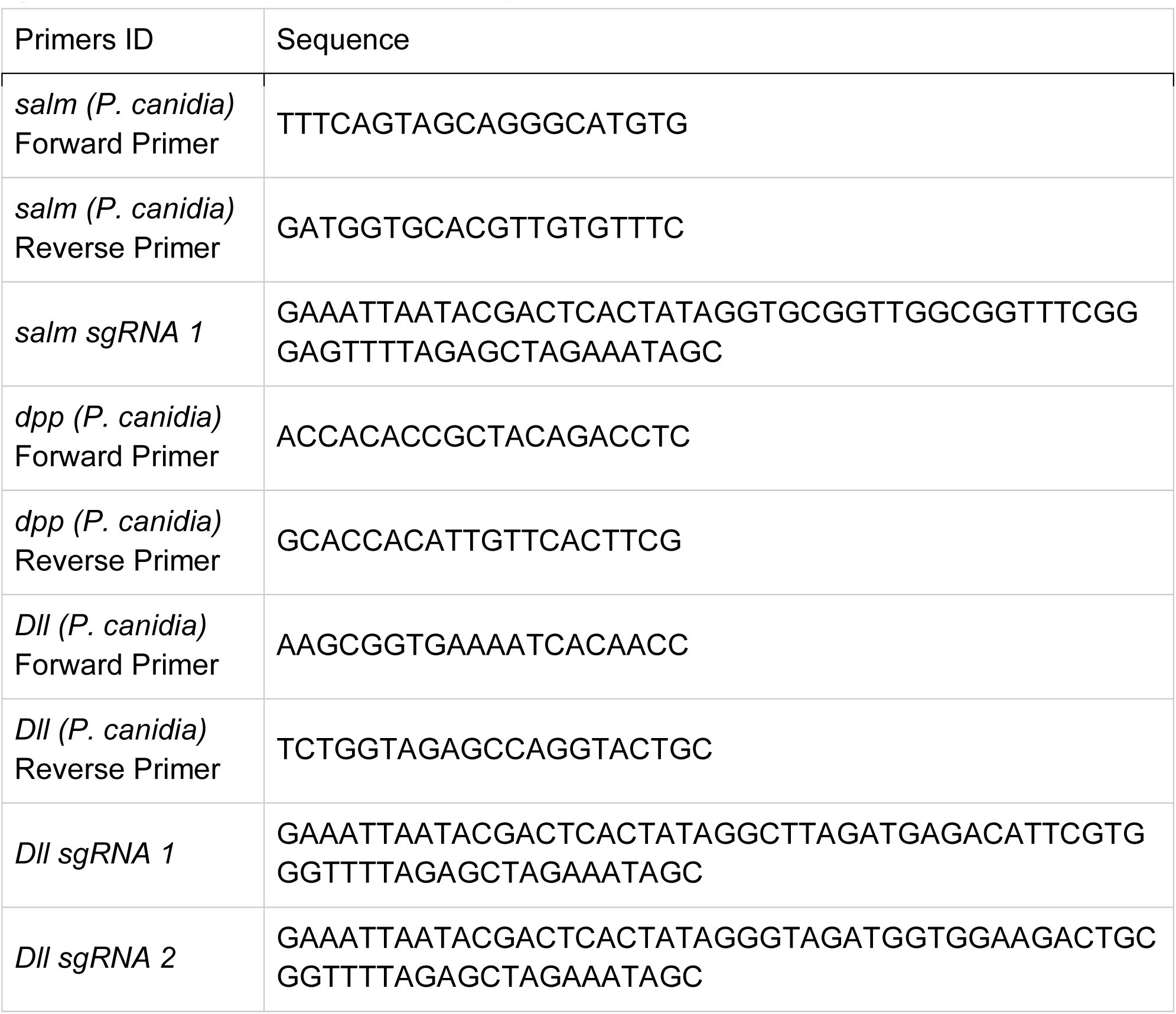
A summary of all the primers used for *in-situ* hybridisations and CRISPR/Cas9 experiments.

**Supplemental Table 2.** Composition of buffers used in immunostaining reactions.

|  |  |
| --- | --- |
| **10X Phosphate-buffered saline (PBS) (In 500mL)** |  |
| Dipotassium hydrogen phosphate ($K_2HPO_4$) | 5.34 g |
| Potassium dihydrogen phosphate ($KH_2PO_4$) | 2.64 g |
| Sodium chloride (NaCl) | 40.9 g |
| Milli-Q Water | To 500 mL |
| **Fix Buffer (In 30 mL)** |  |
| 500 mM PIPES ($C_8H_{18}N_2O_6S_2$) pH 6.9 | 6 mL |
| 500 mM EGTA ($C_{14}H_{24}N_2O_{10}$) pH 6.9 | 60 μL |
| 20% Triton™ X-100 | 1.5 mL |
| 1 M Magnesium sulfate ($MgSO_4$) | 60 μL |
| Milli-Q Water | 22.4 mL |
| 37% Formaldehyde ($CH_2O$) | Add 55 μL per 500 μL of Fix Buffer |
| **Block Buffer (In 40 mL)** |  |
| 1 M Tris-HCl pH 6.8 | 2 mL |
| 5 M Sodium chloride (NaCl) | 1.2 mL |
| 5 mg/mL Bovine Serum Albumin (BSA) | 0.2 g |
| Milli-Q Water | 35.8 mL |
| **Wash Buffer (In 200 mL)** |  |
| 1 M Tris-HCl pH 6.8 | 10 mL |
| 5 M Sodium chloride (NaCl) | 6 mL |
| 20% IGEPAL-CA630 | 5 mL |
| 1 mg/mL Bovine Serum Albumin (BSA) | 0.2 g |
| Milli-Q Water | 179 mL |
| **Mounting media** |  |
| Tris-HCl pH 8.0 | 20 mM |
| N-propyl gallate | 0.5% |
| Glycerol | 60% |

**Supplemental Table 3.** Composition of buffers used in *in-situ* hybridisation reactions.

|  |  |
| --- | --- |
| **10X Phosphate-buffered saline (PBS) (In 500 mL)** |  |
| Dipotassium hydrogen phosphate ($K_2HPO_4$) | 5.34 g |
| Potassium dihydrogen phosphate ($KH_2PO_4$) | 2.64 g |
| Sodium chloride (NaCl) | 40.9 g |
| RNase-free water | To 500 mL |
| **1X Phosphate-Buffered Saline, 0.1% Tween® 20 Detergent (PBST) (In 50 mL)** |  |
| 1X Phosphate-buffered saline (PBS) | 50 mL |
| Tween® 20 | 50 µL |
| **20X Saline Sodium Citrate Buffer (SSC) (In 1000 mL)** |  |
| 3 M Sodium Citrate (NaCl) | 175.3 g |
| Trisodium citrate ($Na_3C_6H_5O_7$) | 88.2 g |
| RNase-free water | 800 mL |
| Use 1 M Hydrochloric acid (HCl) to adjust pH to 7.0<br>Make up volume of buffer to 1 L using RNase-free water<br>Autoclave the buffer before use |  |
| **Pre-hybridization buffer (In 40 mL)** |  |
| Formamide ($CH_3NO$) | 20 mL |
| 20X Saline Sodium Citrate Buffer (SSC) | 10 mL |
| Tween® 20 | 40 µL |
| RNase-free water | 10 mL |
| **Hybridisation Buffer (In 40 mL)** |  |
| Formamide ($CH_3NO$) | 20 mL |
| 20X Saline Sodium Citrate Buffer (SSC) | 10 mL |
| Tween® 20 | 40 µL |
| Salmon Sperm DNA | 40 µL |
| Glycine ($C_2H_5NO_2$) (100 mg/mL) | 40 µL |
| RNase-free water | 10 mL |
| **Block Buffer (In 50 mL)** |  |
| 1X Phosphate-buffered saline (PBS) | 50 mL |
| Tween® 20 | 50 µL |
| Bovine Serum Albumin (BSA) | 0.1 g |
| **Alkaline phosphatase buffer (In 20 mL)** |  |
| Tris Hydrochloride (Tris-HCl) (pH 8.0) | 2 mL |
| 5 M Sodium chloride (NaCl) | 400 µL |
| 200 mM Magnesium chloride (MgCl<sub>2</sub>) | 250 µL |
| Tween® 20 | 20 µL |
| RNase-free water | To 20 mL |

**Supplemental Table 4.**
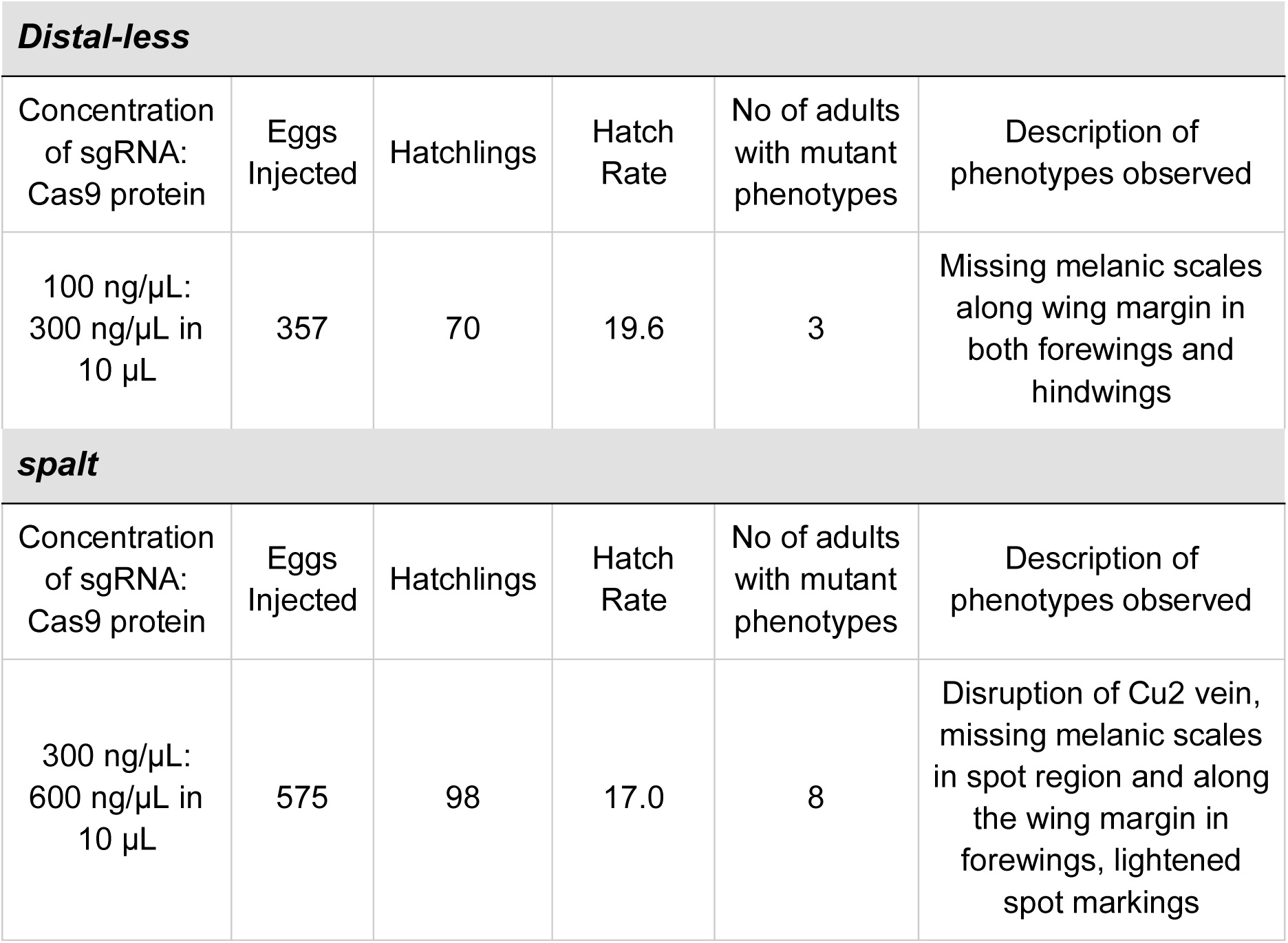
Summary of CRISPR/Cas9 experiments for *Distal-less* and *spalt* knockouts.

| ***Distal-less*** |  |  |  |  |  |
| --- | --- | --- | --- | --- | --- |
| Concentration of sgRNA:<br>Cas9 protein | Eggs Injected | Hatchlings | Hatch Rate | No of adults with mutant phenotypes | Description of phenotypes observed |
| 100 ng/μL:<br>300 ng/μL in<br>10 μL | 357 | 70 | 19.6 | 3 | Missing melanic scales along wing margin in both forewings and hindwings |
| ***spalt*** |  |  |  |  |  |
| Concentration of sgRNA:<br>Cas9 protein | Eggs Injected | Hatchlings | Hatch Rate | No of adults with mutant phenotypes | Description of phenotypes observed |
| 300 ng/μL:<br>600 ng/μL in<br>10 μL | 575 | 98 | 17.0 | 8 | Disruption of Cu2 vein, missing melanic scales in spot region and along the wing margin in forewings, lightened spot markings |

## Probe sequence of *B. anynana dpp* used for in situ hybridisation

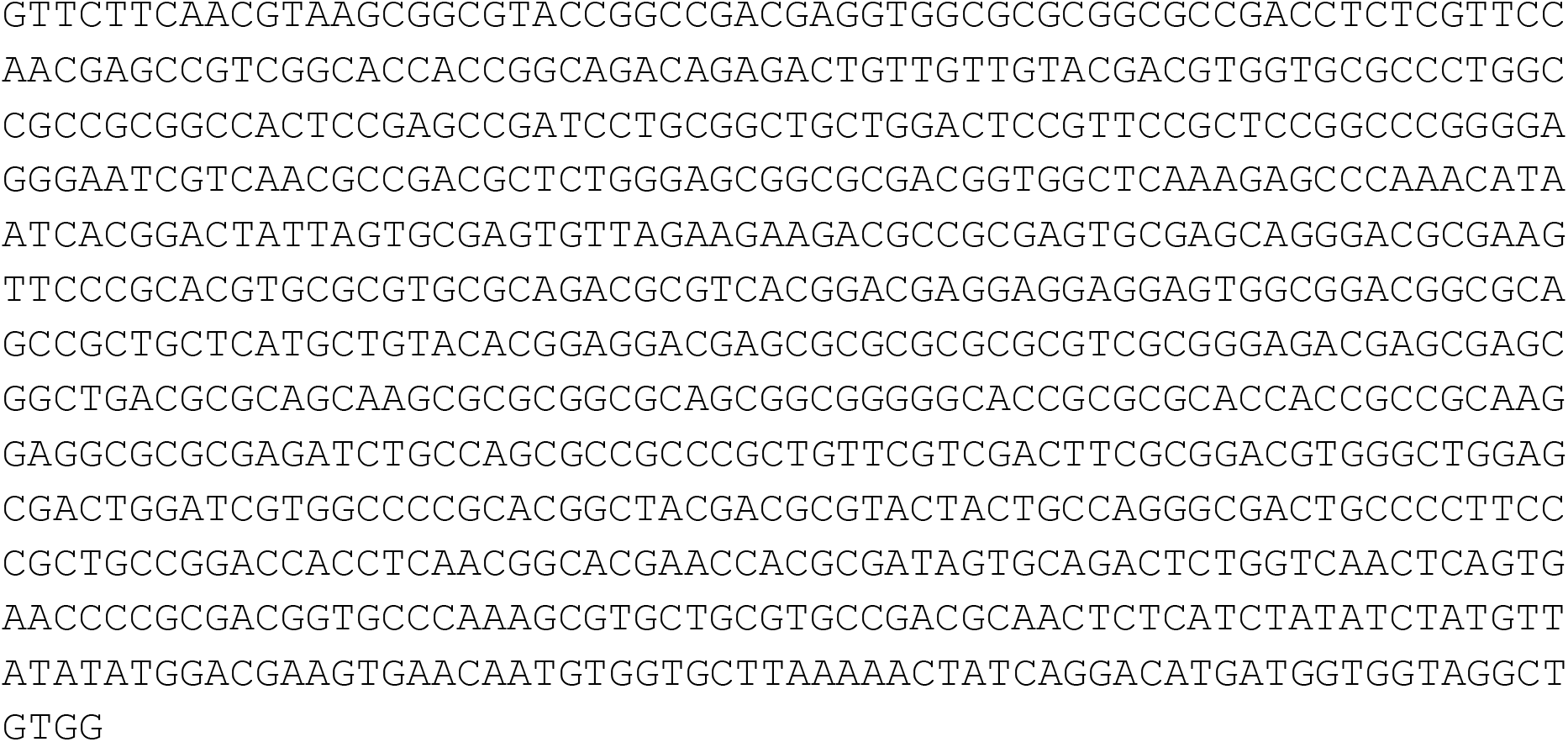

## Probe sequence of *P. canidia dpp* used for in situ hybridisation

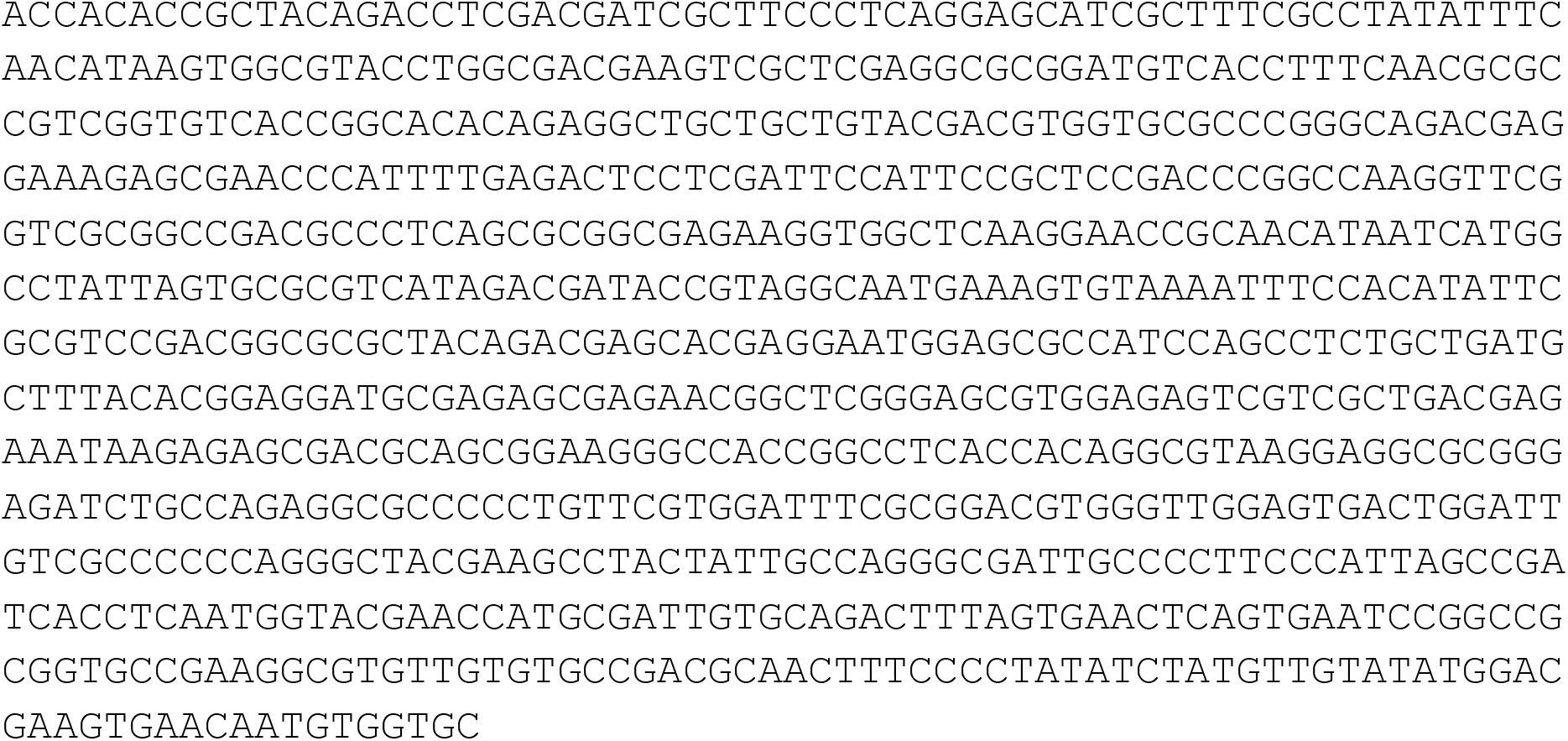

## Sequence of *P. canidia Distal-less* and site of CRISPR targets

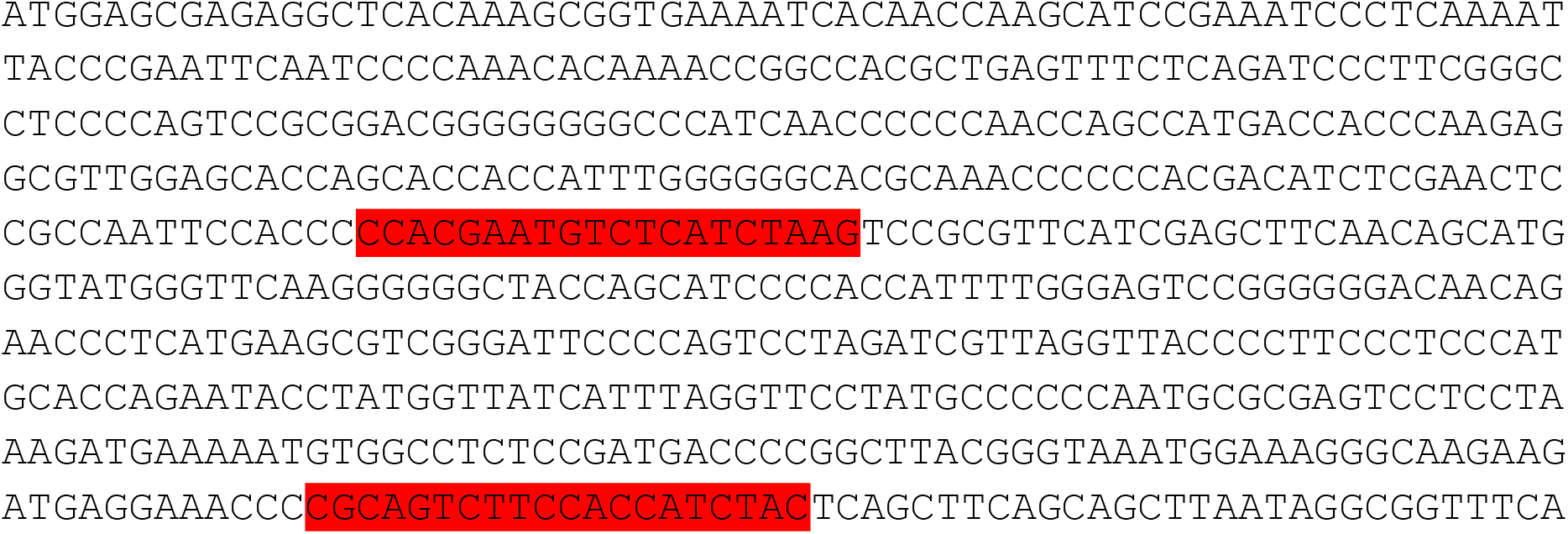
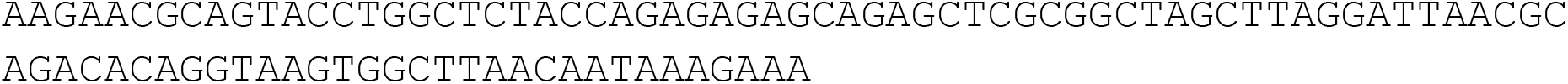

## Sequence of *P. canidia spalt-major* and site of CRISPR targets

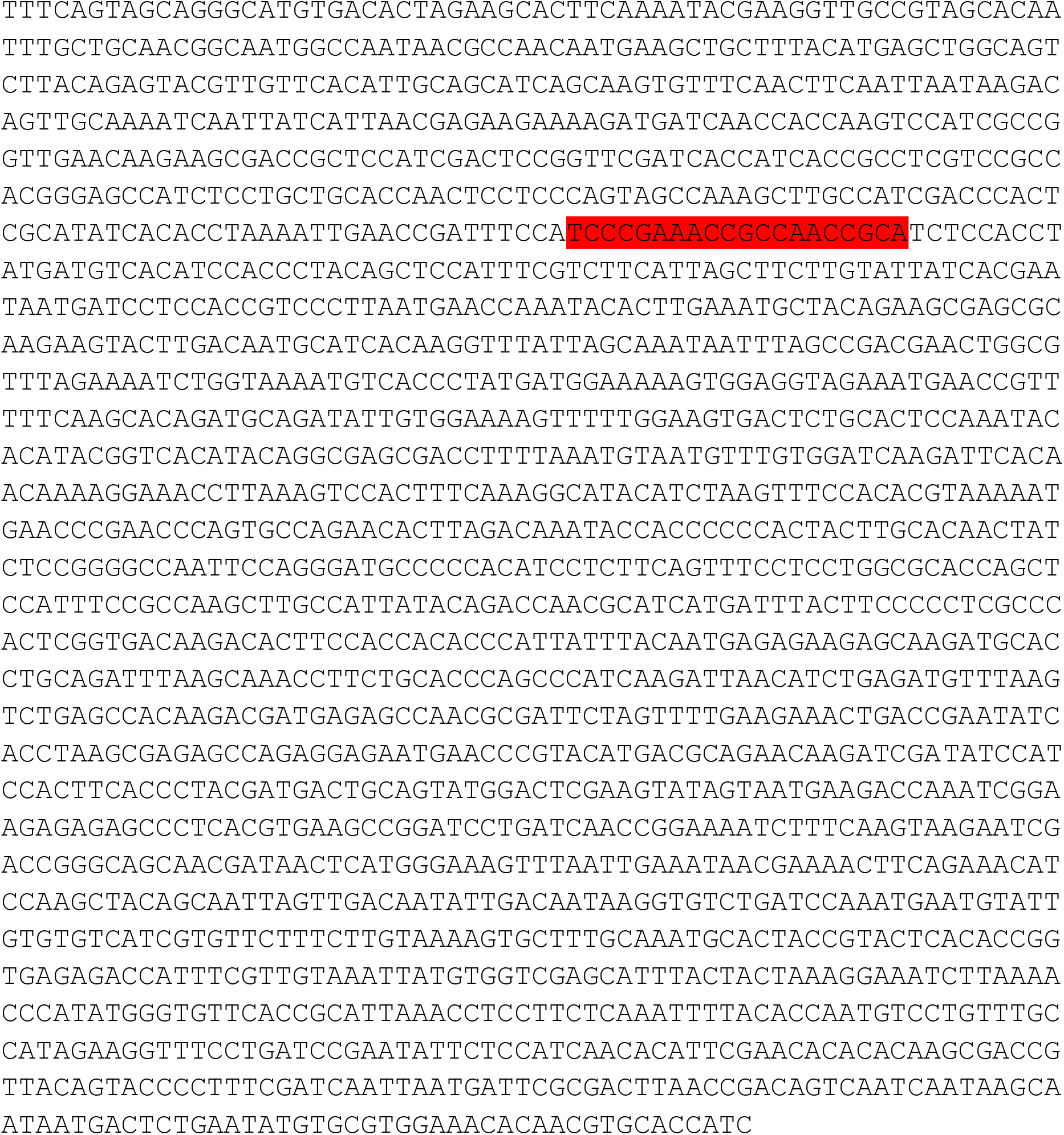

